# Identification and characterization of centromeric sequences in *Xenopus laevis*

**DOI:** 10.1101/2020.06.23.167643

**Authors:** Owen K Smith, Charles Limouse, Kelsey A Fryer, Nicole A Teran, Kousik Sundararajan, Rebecca Heald, Aaron F Straight

**Affiliations:** Department of Biochemistry, Stanford University School of Medicine, 279 Campus Drive, Beckman Center 409, Stanford, CA 94305-5307; Department of Chemical and Systems Biology, Stanford University School of Medicine, Stanford, CA 94305; Department of Genetics, Stanford University School of Medicine, Stanford, CA 94305-5120; Department of Molecular and Cell Biology, University of California Berkeley, 142 Life Sciences Addition #3200, Berkeley, CA 94720-3200

**Keywords:** centromere, *Xenopus laevis*, repetitive sequences, CENP-A, epigenetics

## Abstract

Centromeres play an essential function in cell division by specifying the site of kinetochore formation on each chromosome for mitotic spindle attachment. Centromeres are defined epigenetically by the histone H3 variant CEntromere Protein A (CENP-A). CENP-A nucleosomes maintain the centromere by designating the site for new CENP-A assembly after dilution by replication. Vertebrate centromeres assemble on tandem arrays of repetitive sequences but the function of repeat DNA in centromere formation has been challenging to dissect due to the difficulty in manipulating centromeres in cells. *Xenopus laevis* egg extracts assemble centromeres *in vitro*, providing a system for studying centromeric DNA functions. However, centromeric sequences in *X. laevis* have not been extensively characterized. In this study we combine CENP-A ChIP-seq with a k-mer based analysis approach to identify the *X. laevis* centromere repeat sequences. By *in situ* hybridization we show that *X. laevis* centromeres contain diverse repeat sequences and we map the centromere position on each *X. laevis* chromosome using the distribution of centromere enriched k-mers. Our identification of *X. laevis* centromere sequences enables previously unapproachable centromere genomic studies. Our approach should be broadly applicable for the analysis of centromere and other repetitive sequences in any organism.

## Introduction

Accurate chromosome segregation during cell division requires the centromere, a nucleoprotein complex assembled on each chromosome that is essential for chromosome segregation. Centromeres provide the assembly site for the mitotic kinetochore that mediates microtubule attachment and error correction during mitosis (Foley and Kapoor 2013). Centromeres are defined epigenetically by the histone H3 variant, CENP-A, the presence of which is both necessary and sufficient for centromere formation (Musacchio and Desai 2017). Unlike histone H3.1 nucleosomes, which are assembled as chromosomes replicate in S-phase, CENP-A nucleosomes are replenished after replication during the next G1 phase of the cell cycle. CENP-A nucleosomes in chromatin appear to epigenetically dictate the sites of new CENP-A incorporation, thereby providing a mechanism for self-maintenance (Zasadzińska and Foltz 2017).

In humans, centromeres form on tandem repeats of a ∼171bp DNA sequence termed α-satellite. Each 171bp monomer shares ∼60% sequence homology with other monomers. Tandem arrays of monomers are repeated in blocks of higher order repeats (HORs) resulting in long stretches of virtually identical repeat sequences (McNulty and Sullivan 2018; Rudd, Schueler, and Willard 2003; Willard and Waye 1987). Investigation into the genetic features required to form stable human artificial chromosomes (HACs) identified repetitive α-satellite DNA as a sufficient component for *de novo* centromere formation (Harrington et al. 1997; Ohzeki et al. 2015). These studies demonstrated that repetitive DNA promotes centromere formation in vertebrates.

Perturbing centromere function in cells often leads to cell death, thus cell-free systems using budding yeast and *Xenopus laevis* egg extracts have been invaluable for studying centromere and kinetochore assembly (Ng and Carbon 1987; Hyman et al. 1992; Sorger, Severin, and Hyman 1994; Desai et al. 1997; Akiyoshi et al. 2010; Moree et al. 2011; Guse et al. 2011). Budding yeast centromeres are defined by a single 125bp DNA sequence that is sufficient to recruit much of the centromere and kinetochore in cell extracts. Like human cells, *Xenopus laevis* builds its centromeres on repetitive sequences (Edwards and Murray 2005) and thus *Xenopus* egg extract provides a unique system to study the functions of repetitive DNA in driving centromere formation. A 174 bp centromeric repeat has been previously identified in *Xenopus laevis* by chromatin immunoprecipitation of CENP-A followed by cloning and sequencing (Edwards and Murray 2005). This repeat sequence termed Frog centromere repeat 1 (Fcr1) forms large repetitive arrays and is AT rich, as has been observed for centromeric repeats from other vertebrates (Manuelidis 1978; McDermid et al. 1986; Melters et al. 2013; Sullivan, Chew, and Sullivan 2017). Fcr1 is only detected on 60%-70% of *X. laevis* centromeres suggesting that there must be other sequence elements that comprise *X. laevis* centromeres. *X. laevis* is an allotetraploid species: the genome is composed of two related subgenomes named the long (L) and short (S) based on the length of the homoeologous chromosomes (Session et al. 2016). Whether there is conservation of centromeric repeats within each subgenome or between homoeologous chromosomes remains unknown.

In this study we comprehensively identified and characterized CENP-A associated sequences in *X. laevis*. Using chromatin immunoprecipitation followed by high throughput sequencing (ChIP-seq) we discovered CENP-A associated DNA sequences. We utilized a k-mer based method that does not depend on an assembled reference genome and thus provides an unbiased approach to identifying sequence motifs present at the centromere. We show that a family of Fcr1 related sequences populate *X. laevis* centromeres and colocalize with the centromere protein CENP-C. Our results demonstrate the sequence diversity at active *X. laevis* centromeres and enable future studies of the function of repetitive elements in centromere formation and function.

## Results

### CENP-A associated sequences are composed of diverse, but related repetitive sequences

To characterize centromeric sequences in *Xenopus laevis* we prepared sequencing libraries from solubilized mononucleosomal fractions of total genomic DNA, henceforth referred to as “input DNA”, as well as from CENP-A and H4 immunoprecipitations. Because *X. laevis* centromeric DNA is repetitive (Edwards and Murray 2005), we performed an alignment-independent analysis based on k-mer counting (Hayden and Willard 2012). We identified repeat sequences enriched for CENP-A by generating 25bp long k-mers from each read set and compared the k-mer composition of the CENP-A immunoprecipitate to the input fraction (Fig S1A). For each k-mer, we calculated a centromere enrichment score by dividing the number of times this k-mer was found in the CENP-A ChIP data by the number of times it was found in the input data, normalized to the size of the sequencing libraries. We identified k-mers that were: i) abundant in the CENP-A ChIP and input samples (found at least 1000 times) and ii) enriched in the CENP-A immunoprecipitate compared to the input based on the centromere enrichment score. This approach identified a population of k-mers that are more prevalent in the CENP-A ChIP than input DNA (Fig 1A). Similar analysis performed with different k-mer lengths yielded the same conclusions (Fig S1B-D).

**Figure 1:**
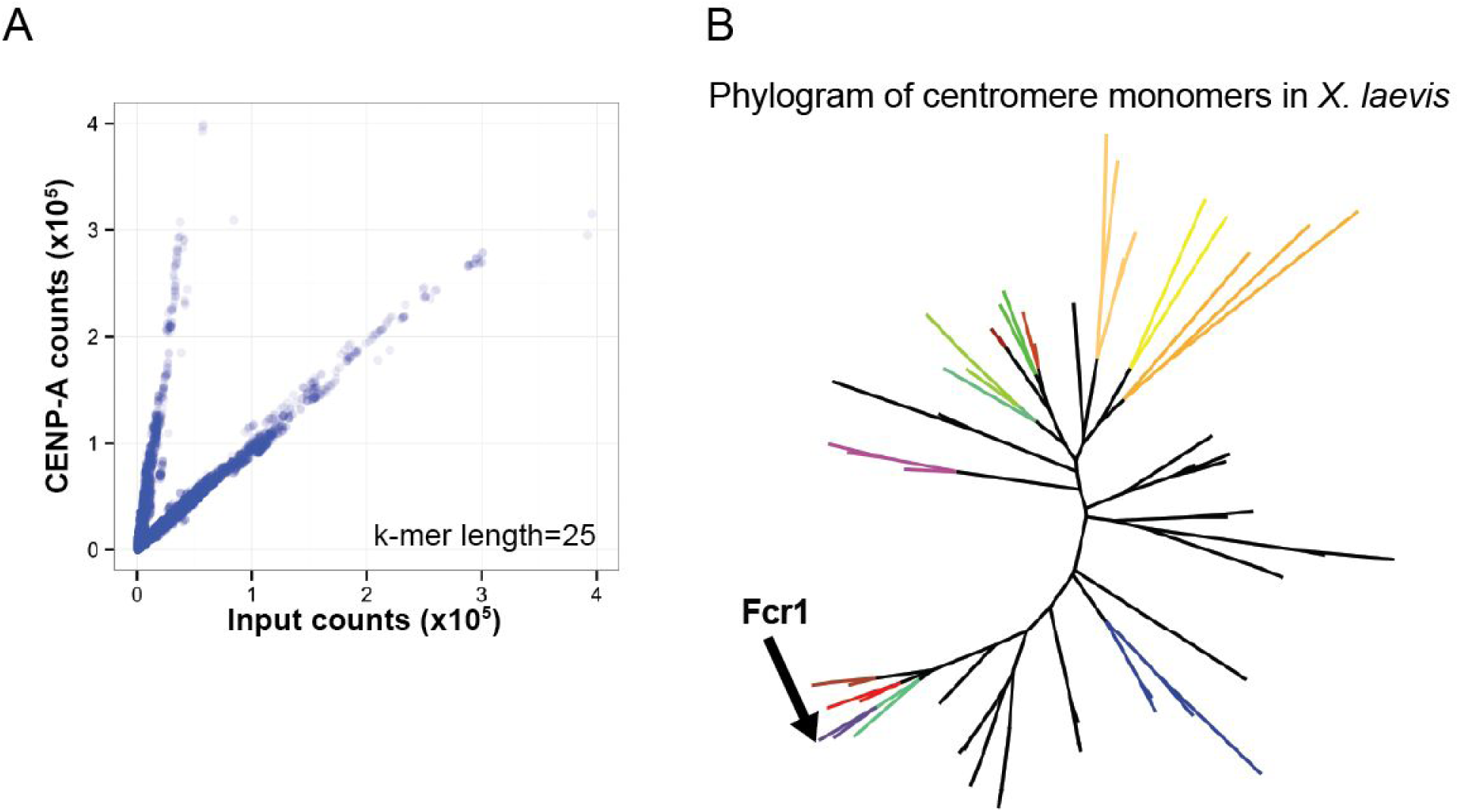
Identification of CENP-A associated sequences by k-mer analysis. A) Scatter plot of 25bp k-mer counts normalized to sequencing depth found in input and CENP-A ChIP-seq libraries. B) Phylogram of representative CENP-A associated sequences that contained a minimum of 80 enriched 25bp k-mers identified as most abundant after clustering by sequence similarity. FCR monomers chosen for FISH experiments are colored and Fcr1 identified by Edwards and Murray is labelled.

**Supplemental Figure 1:**
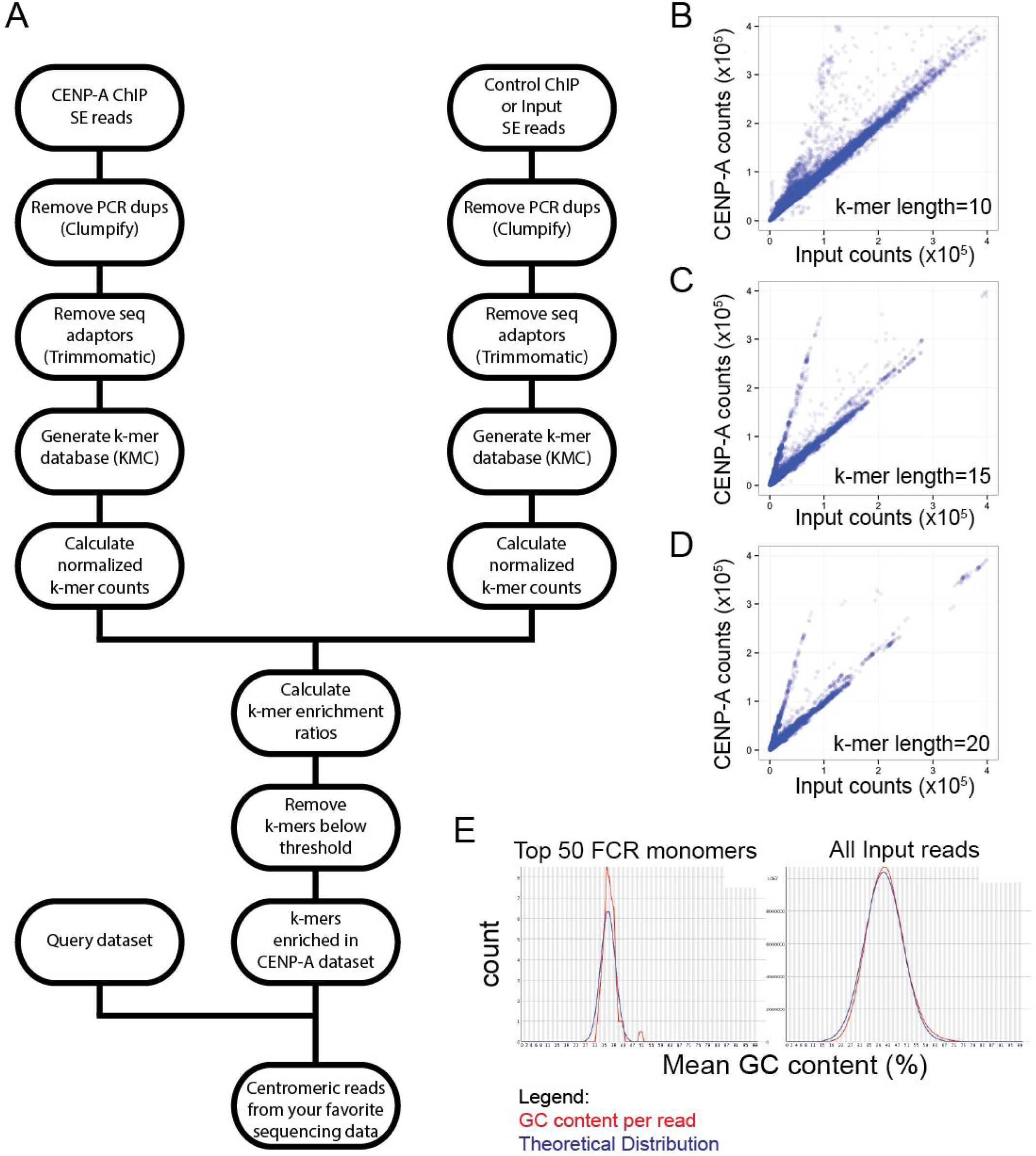
A) Overview of pipeline to identify centromeric k-mers and sequences that contain centromeric k-mers. There are three inputs. The two at the top are reads that will be compared by k-mer content (i.e a ChIP dataset vs an input dataset). The third input, on the left, is a set of sequences that will be sorted by presence of centromeric k-mers. B, C, D) Scatter plots similar to Figure 1A. Scatter plots of normalized k-mer count in CENP-A ChIP (y-axis) vs input (x-axis) for k-mer lengths 10bp (B), 15bp (C), and 20bp (D). E) Histogram of GC-content for top 50 FCR monomers used to generate phylogram in Figure 1B (top). Histogram of GC-content for all input reads (bottom). GC-count per read shown in red and theoretical GC content distribution shown in blue. Plot generated using FASTQC.

To understand the diversity of centromeric sequences in *X. laevis* we isolated reads from our CENP-A ChIP-seq libraries that contained at least one CENP-A enriched k-mer. We then hierarchically clustered these CENP-A ChIP-seq reads by sequence similarity to generate a representative sequence for each cluster. A phylogram of these enriched 150bp representative sequences illustrates the diversity of CENP-A associated repeat sequences (Fig 1B). The previously identified Fcr1 sequence (Edwards and Murray 2005) was present on one clade of the tree validating both the experimental approach and the k-mer based analysis used to identify repetitive elements. All of these sequences were homologous to Fcr1, thus we refer to these as frog centromere repeat (FCR) monomers. The FCR monomer sequences are 150bp long due to the sequencing length and MNase digestion, and thus represent the core position of the dyad axis of the CENP-A nucleosome on the ∼174bp long centromeric monomer. On average the FCR monomers were 39.5% GC (Fig S1E), similar to Fcr1 (Edwards and Murray 2005). Overall, our data demonstrate that the repetitive sequences associated with *X. laevis* CENP-A are related to the previously identified Fcr1, but that they comprise a diverse, previously uncharacterized family.

### FCR monomers vary in their abundance and chromosome specific localization

To validate the centromeric localization of these sequences we performed fluorescence *in situ* hybridization (FISH) combined with immunofluorescence for the constitutive centromere associated protein, CENP-C. Using *X. laevis* sperm nuclei incubated in *X. laevis* egg extract we identified the percentage of centromeres to which each FCR monomer localized (Fig 2A). We confirmed that the previously identified Fcr1 sequence, a member of FCR monomer 16 subfamily, localized to approximately 60% of centromeres (Edwards and Murray 2005). Several other FCR monomers also displayed a similar localization to 60% of centromeres. However, some FCR monomers were present at fewer centromeres, suggesting that these may be FCR variant sequences that are specific to a subset of chromosomes (Fig 2A). Most FCR monomer FISH signals were centromeric with some localization expanded beyond the CENP-C signal, but with very little off-centromere localization, indicating that FCR monomers are found almost exclusively in centromere-associated regions of chromosomes.

**Figure 2:**
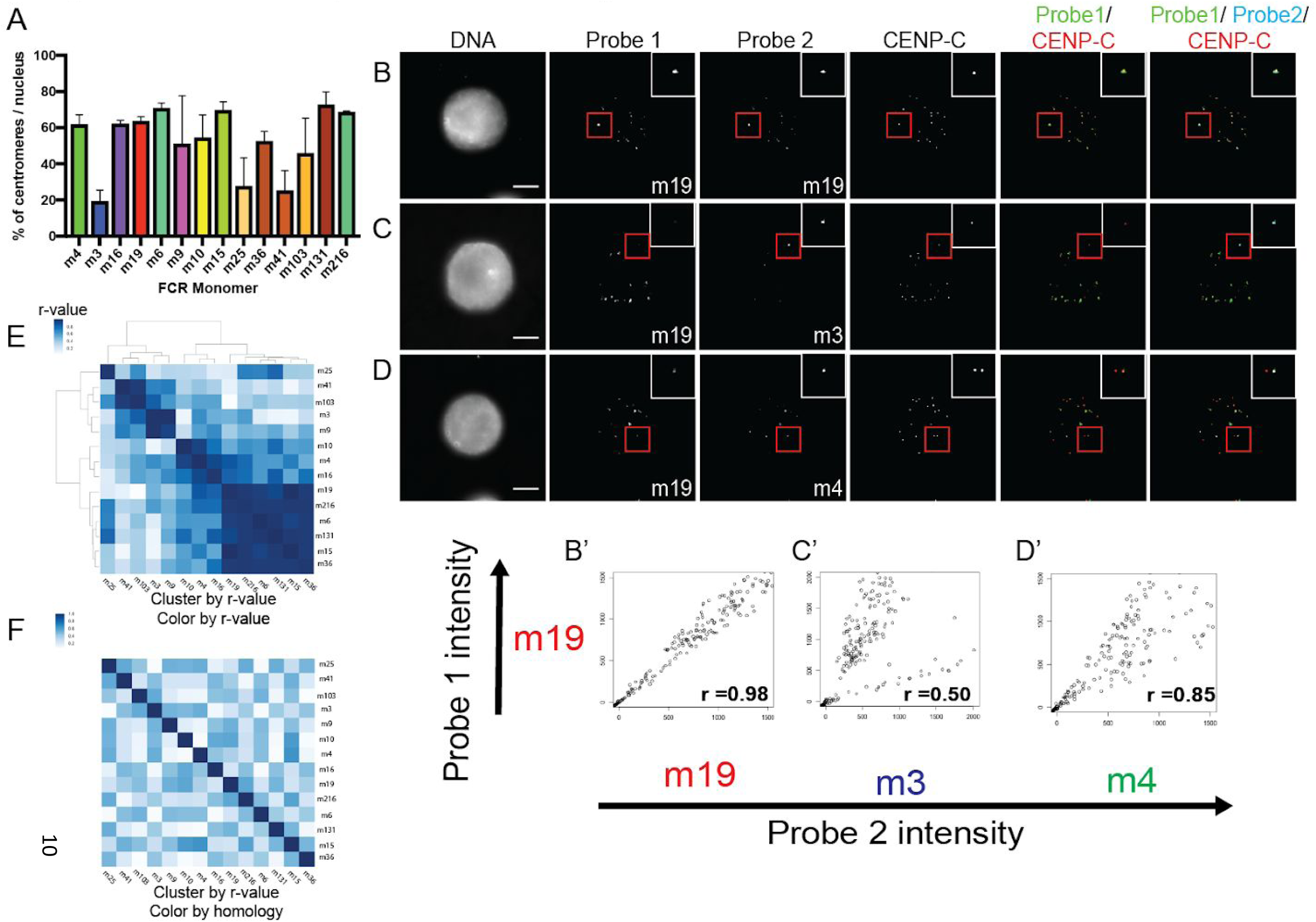
FCR monomers exhibit distinct centromeric localization independent of sequence similarity. A) Bar plot of the percentage of centromeres per nucleus that are positive for a given FCR monomer. Bar color corresponds to color on phylogram. Averages of two independent experiments are shown with standard error displayed. B,C,D) Maximum projection images of two-color FISH with immunofluorescence for the centromere marker, CENP-C. B) FCR monomer 19 vs FCR monomer 19, C) FCR monomer 19 vs FCR monomer 3, D) FCR monomer 19 vs FCR monomer 4. Scale bar = 10µM. B’, C’, D’) Scatter plots of background subtracted probe intensities for each centromere from two-color FISH experiments. Pearson coefficients are displayed in the top left corner. E) Clustered heatmap of FCR monomer Pearson correlation to other FCR monomers as determined by two-color FISH. F) Heatmap ordered based on FISH Pearson correlation clustering, color map displays sequence similarity between FCR monomers.

**Supplemental Figure 2:**
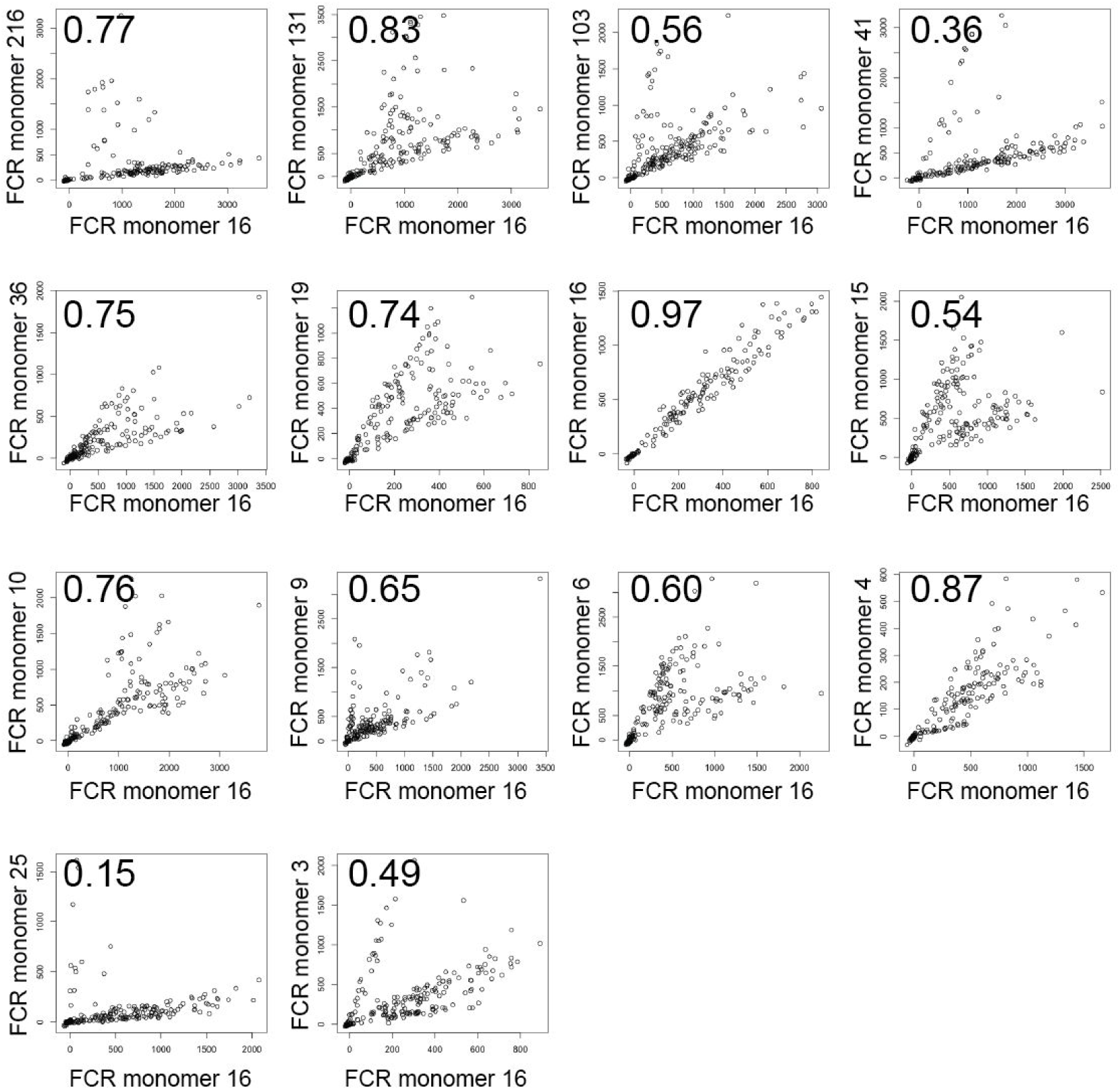
Scatter plots similar to Figure 2A’,B’,C’. Scatter plots of background subtracted probe intensities for each centromere from two-color FISH experiments. Pearson coefficients are displayed in the top left corner. These scatter plots have FCR monomer 16 on the x-axis compared to other FCR monomers tested on the y-axis.

We hypothesized that the distinct branches on the phylogram of CENP-A associated sequences (Fig 1B) might have originated from centromeric sequences found on the parental subgenomes that gave rise to the allotetraploid *Xenopus laevis* genome. To test this hypothesis, we performed two-color FISH combined with immunofluorescence in order to determine whether the different FCR monomers are present on distinct sets of chromosomes. We performed pairwise localization for 14 FCR monomers. Labelling the same FCR monomer sequence in two different colors resulted in overlapping FISH signals with similar intensities (Fig 2B, B’). In contrast, colocalization of different FCR monomers resulted in variable localization and intensities. Some probes appeared to be mutually exclusive such that if a centromere was positive for one probe it was rarely positive for the other (Fig 2C, C’). These likely correspond to FCR monomers occupying distinct chromosomes. Other pairs of FCR monomers were observed on a common set of chromosomes but to a different degree (Fig 2D, D’). In these cases, some chromosomes appeared positive for both probes but with a stronger signal for one of them. Additionally, some FCR monomers appeared equally abundant on individual centromeres (Fig S2). To investigate the degree to which different FCR monomers colocalized, we calculated Pearson correlation coefficients for the intensity of each set of probes at each centromere (Fig 2B’, C’, D’ bottom right) and clustered these correlations. We observed a set of highly correlated probes (Fig 2E) all of which are found at approximately 60% of centromeres (Fig 2A). Therefore, FCR monomers that localize to the majority of chromosomes are likely to be found together on the same centromeres.

To test if the colocalization of different monomers was due to sequence similarity between monomers or due to the localization of distinct monomers on the same chromosome, we colored the heatmap of FISH probes by sequence similarity, but maintained the clustering by colocalization (Fig 2F). This resulted in a loss of the clustered structure indicating that FCR monomers that colocalize are not necessarily closely related at the sequence level. The lack of sequence similarity among FCR monomers that were correlated by FISH indicates that the branches on the phylogram do not predict colocalization. Thus, pairwise FISH analysis of related sequences that all share similarity with the originally identified Fcr1 reveals that not all FCR monomers are found on the same number of chromosomes and that different chromosomes have distinct centromeric repetitive arrays.

### Identification of centromeric repeat arrays on each *X. laevis* chromosome

We next identified the location of the centromeres on each chromosome by identifying the regions in the genome that contained the most CENP-A associated k-mers. The histogram of centromere enrichment scores has a median near one with a long tail of enrichment values above 1 (Fig 3A). Using an updated version of the *X. laevis* genome that has a single contig per chromosome (X. laevis genome v10.2) we partitioned each chromosome into non-overlapping 50kb segments and identified the 50kb segments that contained CENP-A enriched k-mers. We selected k-mers for alignment to the genome by increasing the threshold for inclusion based on the magnitude of the centromere enrichment scores. At low enrichment scores we observed that most genome segments contained at least one k-mer (Fig 3A, B). Increasing the centromere enrichment scores we used as a cutoff resulted in a steady decrease in the percentage of genome segments containing an enriched k-mer (Fig 3B). Upon increasing the threshold for inclusion from an enrichment value of 3.31 to 3.46, the number of genome segments containing an enriched k-mer dropped from 77.67% of all segments to 0.15% of segments. This dramatic reduction in genome segments containing enriched k-mers arose from a modest reduction in the total number of k-mers from 3,825 to 3,350 enriched k-mers. Further increasing the stringency caused no change in the percentage of genome segments with enriched k-mers, yielding 84 50kb segments that represent the centromere repeat containing segments of the *X. laevis* genome.

**Figure 3:**
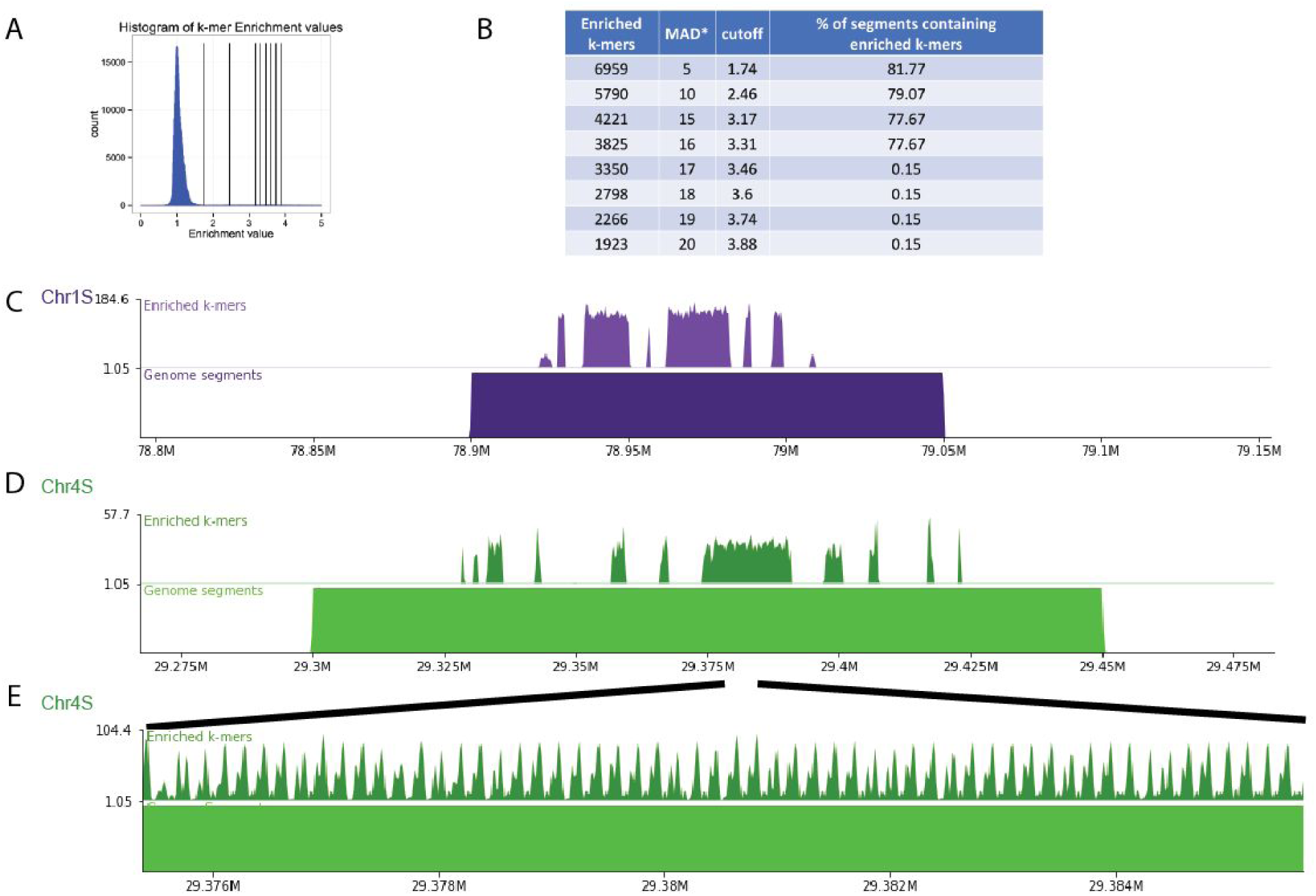
Identification of centromeres on *Xenopus laevis* chromosomes. A) Histogram of centromere enrichment scores for 25bp k-mers. Enrichment scores are the ratio of normalized k-mer counts for the CENP-A dataset over the input dataset. Vertical lines display stringency cutoffs of (1, 2, 5, 10, 15, and 20) median absolute deviations away from the median enrichment value. B) Table displaying the number of enriched 25 bp k-mers, the median absolute deviations (MAD*) away from the median used as the cutoff value, the enrichment value cutoff, and the percentage of genome segments containing an enriched k-mer. C, D, E) Representative genome browser images with aligned enriched k-mers (top) and aligned genome segments (bottom). E is a zoom in on a region in D.

**Supplemental Figure 3:**
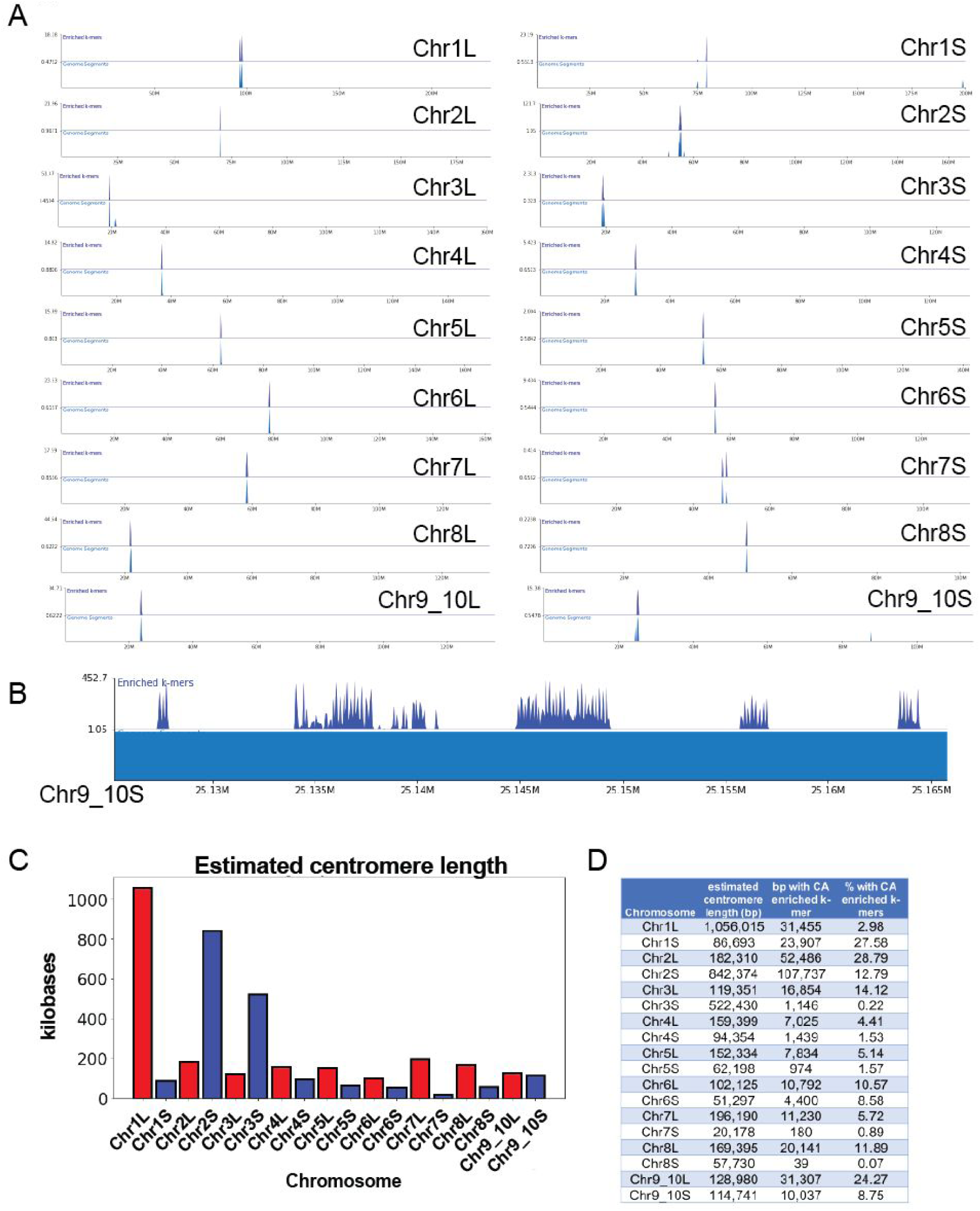
A) Chromosome overviews showing the entire scaffold for each chromosome in *X. laevis* genome. Top track is alignment of enriched 25bp k-mers, defined as 17 Median Absolute Deviations above the median of all CENP-A/Input enrichment values. Bottom track is 50kb genome segments that contain enriched k-mers. Most chromosomes have one location where enriched k-mers map. B) Zoom in of the genome browser view of the centromeric region on Chromosome 9_10S showing that arrays of repetitive regions identified in this study are interspersed with other sequences. These intervening sequences could be repetitive or non-repetitive, but do not contain CENP-A enriched k-mers. C) Barplot of centromere size on each chromosome in kilobases. Exact size of the centromeric repetitive array is estimated by identifying the distance between the first and last base pairs that have a CENP-A enriched k-mer aligned per chromosome. These estimates include the intervening sequences between centromeric repetitive arrays shown in Fig S3B, D) Chart of the centromere length (as defined in C), base pairs with CENP-A enriched k-mers, and percentage of centromere that contain CENP-A enriched k-mers for each chromosome.

We mapped these 84 segments onto the *X. laevis* genome and found that almost all chromosomes contained a single locus of centromere repetitive arrays (Fig S3A). On each chromosome the location of 50kb genome segments with CENP-A enriched k-mers were frequently contiguous (Fig 3C, D). Within individual 50kb segments we found gaps between regions containing CENP-A enriched k-mers (Fig S3B). Both of these observations are consistent across chromosomes, including the Chr 9_10 homoeologous pair, which arose from a chromosome fusion event. In human centromeres, highly homogeneous repetitive arrays at the core of the centromere become less homogeneous in flanking genomic regions (Miga et al. 2014). Similarly, when we examined the edges of the repetitive array we observed a lower density of enriched k-mers that were less contiguous (Fig 3C, D). Local alignment of the enriched k-mers in each 50 kb segment revealed peaks and valleys that span the genome segments (Fig 3E). Interestingly, the distance between peaks is ∼170 bp, similar to the monomer size of human centromeric repeats and the size of the originally reported Fcr1 (174bp). Not all monomer sequences in the repetitive array possessed k-mer peaks of an identical shape, further suggesting that these tandem arrays are not composed of identical repeated monomers, but of diverse monomers that potentially form higher order repeats. We estimate that the size of the repetitive array on each chromosome can vary from as little as ∼20kb on Chr 7S up to ∼1Mb on Chr 1L (Fig S3C). These estimates are based on the distance between the first and last base pair where a CENP-A enriched k-mer aligned after manual selection of the centromeric region based on 50kb genome segments. These estimates include the gaps between repetitive arrays on each chromosome. It is important to note that centromeres based on these estimates are only partially covered by CENP-A enriched k-mers (Fig S3D). Whether these gaps also contain CENP-A nucleosomes, but were not identified by this k-mer analysis because the sequences are not repetitive is unknown. It is possible that these centromere length calculations are underestimates because repetitive array lengths on genome assemblies could be limited by the long read sequencing data from which they are made. However, these centromere size estimates are similar to CENP-A containing arrays observed in humans (Miga et al., 2014; Sullivan et al., 2011).

### Chromosome specific assignment of centromere sequences

To determine which k-mers were present in each centromeric region we clustered each 50kb genome region by the similarity of their CENP-A enriched k-mers (Fig 4A). We observed that some groups of k-mers were only found on genome segments from the same chromosome (Chr 4S), while other groups of k-mers, seen as vertical stripes on the heatmap, were found on genomic regions from several different chromosomes. Surprisingly, no strong correlations could be made for subsets of k-mers localizing to one ancestral subgenome or the other. To illustrate the relationship between the k-mer content on the two subgenomes we reordered the genome segments (rows) by their chromosome of origin while maintaining the clustering by k-mer similarity (Fig 4B). This shows that some homoeologous chromosomes can have very similar k-mer spectra (e.g. Chr 1L and 1S), while other homoeologous chromosome pairs have a distinct centromeric makeup (Chr 2, Chr 4, and Chr 9_10 pairs) (Fig 4B).

**Figure 4:**
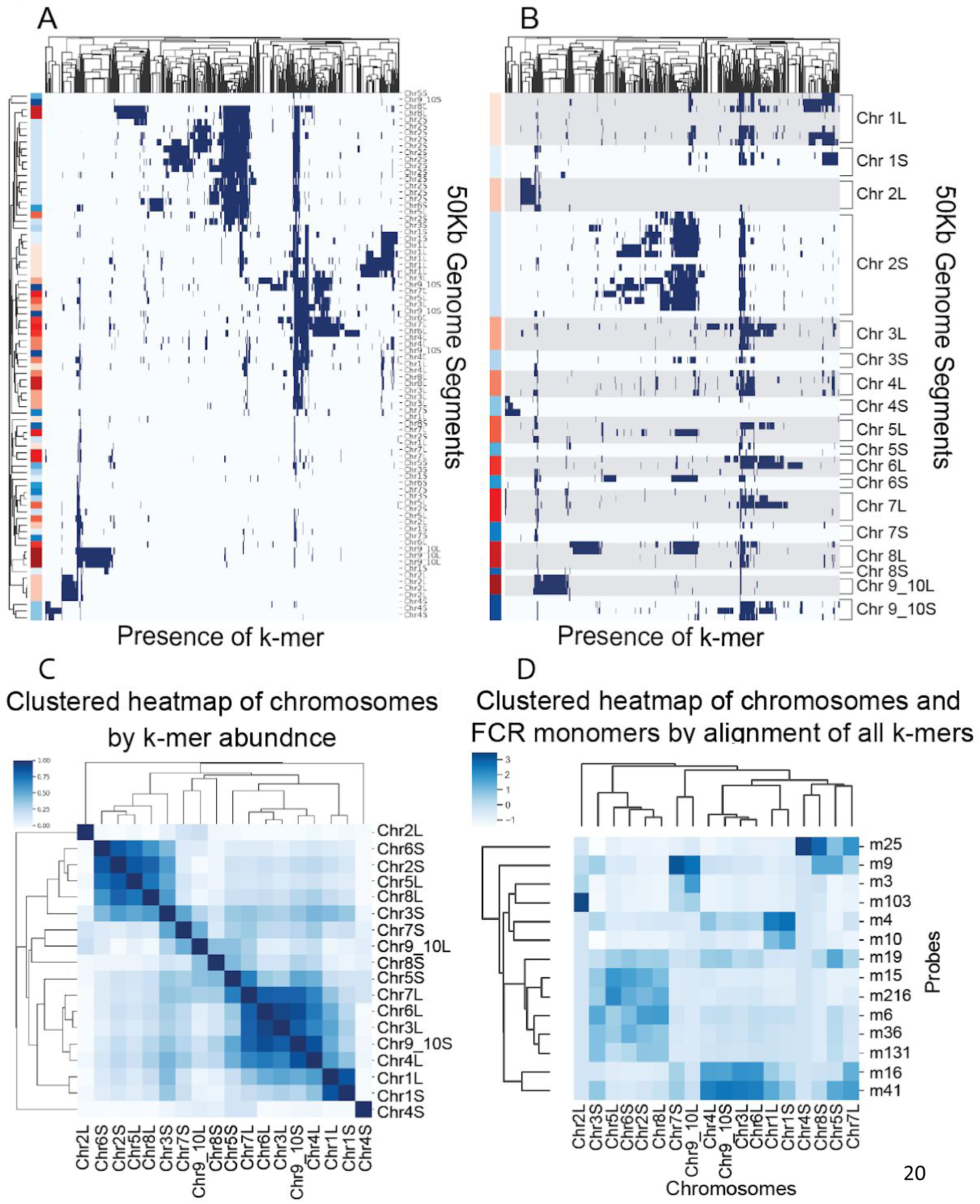
Assignment of FCR monomers to chromosomes by k-mer content. A) Clustered heatmap showing the presence (blue) or absence (white) of individual enriched k-mers on each centromeric genome segment. Both rows and columns are clustered to show k-mers and segments that display similar distributions. Genome segments, on the y-axis, are labelled on the left side indicating the L subgenome (blue), S subgenome (red). B) Similar to A, but the genome segments, y-axis, are not ordered based on similar k-mer content, and are instead listed by chromosome. L subgenome chromosomes are shaded with grey for clarity. C) Clustered heatmap of chromosomes by abundance of CENP-A enriched k-mers. By combining 50kb genome segments from each chromosome, an array of counts for each k-mer was used to generate an Euclidean distance between chromosomes used for clustering. Coloring of heatmap is (1-Euclidean distance). D) Clustered heatmap of counts reported from bowtie1 of the number of times any k-mer from each FCR monomer aligns to each chromosomal contig.

**Supplemental Figure 4:**
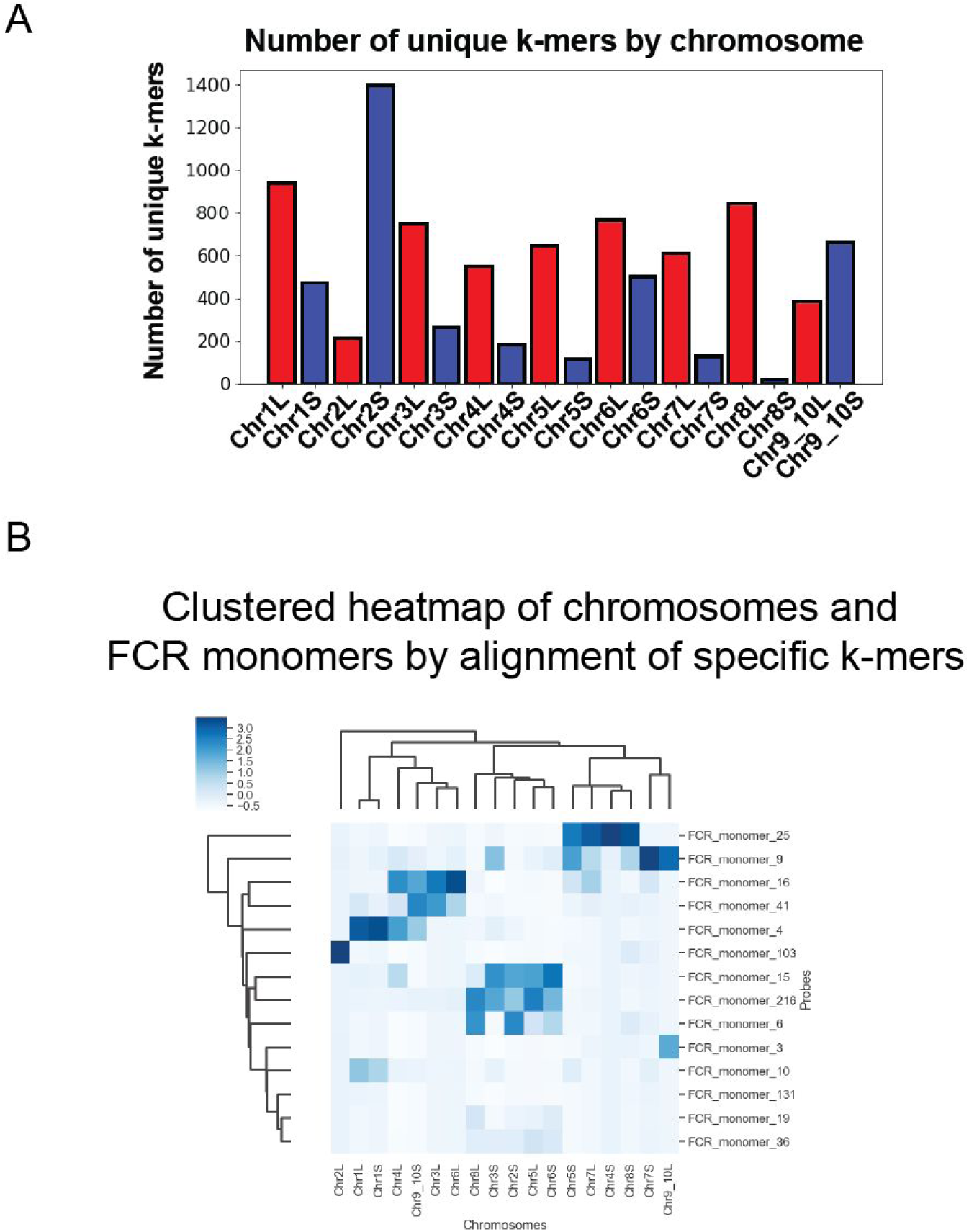
A) Bar plot of the total number of unique k-mers found on each chromosome based on centromeric genome segments. Chromosomes from the L subgenome are shown in red and from the S subgenome are shown in blue. B) Heatmap similar to Figure 4C. Clustered heatmap of counts reported from bowtie1 of the number of times any unique k-mer from each FCR monomer aligns to each chromosomal contig. Alignment counts were normalized by the number of k-mers unique to each FCR monomer.

No clear trends emerged for the similarity of k-mer signatures between homoeologous chromosomes or between chromosomes from the same ancestral genome suggesting a diverse evolutionary history of *Xenopus* centromeres. To evaluate the similarity of centromeres between homoeologous chromosomes we clustered the chromosomes by the frequency with which shared k-mers were found between chromosomes (Fig 4C). Only one homoeologous pair (Chr 1L and Chr 1S) cluster together based on shared k-mer abundance. Chromosomes from both the L and S subgenome cluster together, and in some cases chromosomes with distinct k-mer content do not cluster with any other chromosomes (Chr 2L and Chr 4S). Chromosomes with fewer unique k-mers (Fig S4A), do not neatly cluster with other sets of chromosomes (Chr 7S, Chr 8S. Chr 9_10L). The clustering of these chromosomes may be determined by the absence of shared k-mers rather than the k-mers they contain. L subgenome chromosomes have more unique centromeric k-mers, with the exception of Chr 2 and Chr 9_10 pairs, which may reflect a higher centromeric mutation rate in that subgenome before the allotetraploidization of *X. laevis*.

We have evaluated the k-mer content of each *X. laevis* chromosome, however, to understand which 174 bp monomer sequences make up the centromeric arrays on each chromosome requires mapping the individual monomers to specific chromosomes. To assign FCR monomer sequences to specific chromosomes we extracted k-mers from the monomers that we localized by FISH (Fig 2) and aligned just those k-mers to individual *X. laevis* chromosomes. Using bowtie (B. Langmead et al. 2009) we allowed sequences to align as many times as possible without mismatches. By measuring the number of times k-mers from an FCR monomer aligned to each chromosome we assessed from which chromosome each FCR monomer was likely derived (Fig 4D). Stronger signal in the heatmap indicates the chromosomes to which the FCR monomers would be predicted to have the strongest hybridization. The clustering of FCR monomers in this heatmap is similar to the clustering from the two-color FISH experiments (Fig 2E). Most notably the dominant cluster by FISH containing FCR monomers 216, 131, 36, 19, 15, and 6 is recapitulated by this method. Additionally, by FISH FCR monomer 25 clustered by itself and by this alignment-based analysis FCR monomer 25 k-mers aligned most to Chr 4S, indicating that Chr 4S is the chromosome detected by FCR monomer 25 in the FISH experiment. Chr 8S also has k-mers from FCR monomer 25, but the repetitive array on this chromosome is short, less than 60kb (Fig S3C) which may explain why only one centromere stains strongly for FCR monomer 25 by FISH. Aligning only 25bp k-mers that are unique to each 150bp monomer and not shared between them generated a very similar heatmap to the alignment of all k-mers (Fig S4B), supporting that the differences between FCR monomers drives the observed signal of chromosomal localizations. Our ability to assign specific FCR monomers to individual chromosomes makes genomic analysis of individual chromosomal centromeres in *X. laevis* possible.

## Discussion

In this study, we characterized the active centromeric regions in *Xenopus laevis* using native MNase CENP-A ChIP-seq. We utilized a k-mer based strategy to functionally define the active centromeric repeat DNA on each chromosome in *X. laevis*. The k-mer counting approach we developed can be applied to study any repeats present in a genome using ChIP-seq or analogous datasets. We found that, in *X. laevis* centromeres, the primary sequences associated with CENP-A are a diverse set of repeat sequences related to Fcr1 (Edwards and Murray 2005). We used sequence mapping and *in situ* hybridization to show that groups of these FCR monomers form repeat arrays that can be either 1) unique to individual chromosomes, 2) shared between subsets of chromosomes with different levels of abundance or 3) mutually exclusive when compared between chromosomes. These observations lead to several different models for how centromeric sequences are established and maintained in *X. laevis*. Homoeologous chromosomes that possess similar k-mer content (e.g. Chr 1L and 1S) suggest that the ancestral chromosome before divergence may have contained this same repetitive centromeric array, and that both homoeologs maintained this HOR. Alternatively, some homoeologous chromosomes have distinct centromeric k-mer spectra (e.g. Chr 4L and 4S). As Chr 4L shares k-mers with other chromosomes, including Chr 1L and 1S, an ancestral centromeric repeat may have been shared between these chromosomes. Chr 4S may have once harbored this same ancestral repeat, but acquired a distinct centromeric repeat that became multimerized and fixed over time. Transposable elements or intrachromosomal recombination may also generate diversity observed between pairs of homoeologs and within each subgenome. The presence of diverse sequences at centromeres suggests that multiple sequences have the capacity for retaining CENP-A and maintaining centromeres.

Although diverse sequences compose centromeres in *X. laevis*, these sequences could have common properties that allow them to be competent for centromere establishment. Studies from fission yeast (Ngan and Clarke 1997), fly (X. Sun, Wahlstrom, and Karpen 1997; Xiaoping Sun et al. 2003; Peacock et al. 1974) and human (Hayden et al. 2013), suggest that multiple distinct repetitive units can have the capacity to harbor active centromeres and may have the ability to form centromeres *de novo*. These studies demonstrate the capacity of repetitive DNA to establish an active centromere in eukaryotic systems. While the role of CENP-A in defining centromeres is undisputed, these investigations highlight the potential function of the underlying centromeric DNA in specification and establishment of centromeres.

In humans, alpha satellite DNA can promote CENP-A assembly in part by providing a binding site for the CENP-B protein (Ohzeki et al. 2002). *X. laevis* appears to lack a centromere binding CENP-B homolog, yet sequence features of *X. laevis* centromeric DNA may facilitate centromere function, as has been suggested for other species (Kasinathan and Henikoff 2018). A recent study demonstrated that certain non-repetitive chromosomal fragments contain the ability to retain CENP-A after transient targeting of HJURP to deposit CENP-A (Logsdon et al. 2019) suggesting that some non-alphoid sequences are competent for CENP-A retention. These studies challenge the notion that centromeres are specified purely by epigenetic factors and motivate the investigation of DNA sequence contributions into centromere maintenance in diverse model organisms.

In this study we have characterized *X. laevis* centromeric repeats, enabling dissection of the genetic determinants of *X. laevis* centromeres. By assigning repeat monomers to specific chromosomes we allow further study of *X. laevis* centromeres using genomic techniques. How DNA elements synergize with centromere assembly factors that epigenetically promote CENP-A nucleosome formation is a key question in centromere formation and inheritance. The approach that we apply to *Xenopus laevis* centromeres should be broadly applicable to the study of centromere or other repeat sequences in any organism.

## Methods

### MNase ChIP-seq library preparation

Adult J-strain *Xenopus laevis* were anesthetized and sacrificed before blood was drawn. For each frog 6-8 aliquots of ∼300µl of blood was washed three times by centrifugation for 5 minutes at 1400g with 1ml of Buffer 4 (15mM Sodium Citrate, 150mM NaCl) to prevent clotting. Two additional washes were performed with 1mL each of Buffer 1 (2.5mM EDTA, 0.5M EGTA, 15mM Tris-HCl pH 7.4, 15mM NaCl, 60mM KCl, 15mM Sodium Citrate 0.5mM spermidine, 0.15mM spermine, 340mM Sucrose, supplemented with 0.1mM PMSF). Cells were resuspended in Buffer 1, pooled and dounced with Wheaton pestle B (30-50x). Cells were checked on a hemocytometer to confirm complete lysis of cell membrane and intact nuclei. After lysis, cells were washed two times with Buffer 3 (15mM Tris-HCl pH 7.4, 15mM NaCl, 60mM KCl, 0.5mM spermidine, 0.15mM spermine, 340mM Sucrose, supplemented with 0.1mM PMSF), and resuspended in 500µl Buffer 3. CaCl_2_ was added to 5mM. Chromatin was digested with 300U MNase 30min at room temperature. Digestion was quenched with a final concentration of 2mM EDTA and 1mM EGTA. To lyse nuclei 10% IGEPAL CA-630 was added to final concentration of 0.05%. Samples were incubated on ice for 10min followed by sedimentation of the chromatin at 5min 1500g, 4°C and removal of the supernatant. Chromatin was resuspended in 500µl Buffer 3 supplemented with 200mM NaCl. The chromatin was extracted by overnight rotation at 4°C. The chromatin was pelleted at ∼16,000Xg for 10min at 4°C. The supernatant was collected and a sample was processed to confirm digestion to mostly mononucleosomes. This supernatant is the ChIP input. The input was precleared by rotation at 4°C for 4hrs to overnight with 100µl Protein A dynabeads prewashed with TBST (0.1% Triton X 100). For immunoprecipitation 5µg of antibody (*X. laevis* CENP-A or H4 Abcam ab7311) was coupled to 20µl of protein A dynabeads that had been washed three times with 400µl TBST by rotation at 4°C in a final volume of 200µl TBST. Antibody bound beads were washed three times with TBST and beads were collected. Precleared beads were collected and the precleared input was split evenly between antibody bound beads. A sample was also taken for input library preparation. Samples were rotated overnight at 4°C to bind nucleosomes to beads. After overnight rotation beads were collected and washed three times with 400µl TBST. Beads were then resuspended in 40µl TE (10mM Tris-HCl pH 8.0, 1mM EDTA) supplemented with 0.1% SDS. ProteinaseK was added to 0.25mg/ml (0.5 µl of 20mg/ml) and samples were incubated at 65°C with 850rpm shaking from 4hrs to overnight. Beads were collected and ChIP samples were transferred to a new tube.

Ampure beads were used to isolate the mononucleosomal fraction. Briefly, 1.6x sample volume of beads were mixed with the sample and incubated for 5min room temperature. Beads were collected and supernatant was removed. Beads were washed 2 times with EtOH and allowed to air dry on magnet for 5 min. Beads were then eluted with 27µl 10mM Tris-HCl pH 8.0. ChIP eluates and input were assessed by high sensitivity Qbit and Agilent Bioanalyzer. Sequencing libraries were prepared with the NEBNext Ultra II DNA Library prep kit with up to 1µg of input or ChIP eluate DNA following the manufacturer’s protocol. Two replicates of each sample were sequenced on MiSeq and one replicate on a HiSeq Illumina NGS sequencers.

### DNA FISH and IF protocol

*Xenopus laevis* CSF-arrested egg extract was prepared as described and supplemented with *Xenopus laevis* sperm nuclei. The extracts were released into interphase for 75-90 min before being fixed with 2% Formaldehyde, sedimented onto poly-lysine coated coverslips and processed for immunofluorescence (French 2017). Briefly, Coverslips were washed quickly with PBS, and Antibody Dilution Buffer (AbDil) (150mM NaCl, 20mM Tris-HCl pH 7.4, 0.1% Triton X-100, 2% BSA, 0.1% Sodium Azide), before blocking in AbDil for 30 min. Samples were incubated in primary antibody for 30 min at room temperature (1µg/ml Rabbit anti-*X. laevis* CENP-C antibody diluted in AbDil), washed quickly three times in AbDil, and incubated in secondary antibody (Donkey anti Rabbit-Alexa 647) diluted 1:1000 in Abdil for 30 min at room temperature. Samples were washed again in AbDil three times.

After immunofluorescence, samples were fixed again in 2.5% formaldehyde diluted in PBS for 10 min and washed three times in PBS. Samples were treated with 100µg/ml RNAse A in PBS for 30 min and washed again in PBS for 30 min. Samples were then dehydrated with an ethanol series for 1-2 min in 70%, 80%, 95%, and 100% EtOH before being allowed to air dry for two min. Probes were diluted and denatured at 75°C for 5 min before being spotted onto coverslips. The coverslips were then inverted onto slides, and incubated on a heat block at 80°C for 10 min. Slides were then transferred to a humid chamber and hybridized at 37°C overnight. Coverslips were floated and inverted off of slides with 4X SSC (0.6M NaCl, 60mM Sodium Citrate) and washed with 4X SSC. Coverslips were then washed three times with 2XSSC pre-warmed to 37°C with 50% formamide for 5min each, three times with 2X SSC prewarmed to 37°C for 5min each, one time with room temperature 1X SSC for 10 min, and one time with room temperature 4X SSC for 5 min. After washing, coverslips were stained with 10µg/ml Hoechst-33342 diluted in AbDil for 10 min, washed one time with PBS containing 0.1% Triton X-100, and one time with PBS, before being mounted (20mM Tris-HCl pH 8.8, 0.5% p-Phenylenediamine, 90% Glycerol) on slides and sealed with nail polish.

Imaging was performed on an IX70 Olympus microscope with a DeltaVision system (Applied Precision) a Sedat quad-pass filter set (Semrock) and monochromatic solid-state illuminators, controlled via softWoRx 4.1.0 software (Applied Precision). Images of sperm nuclei were acquired using a 60x 1.4 NA Plan Apochromat oil immersion lens (Olympus). Images were acquired with a charge-coupled device camera (CoolSNAP HQ; Photometrics) and digitized to 16 bits. Z-sections were taken at 0.2-µm intervals. Displayed images of sperm nuclei are maximum intensity projections of z-stacks.

FISH probes were generated using random hexamer priming. To generate FISH probes 150bp FCR monomer sequences were ordered as GeneBlocks from IDT. GeneBlocks were blunt ligated into the pJET1.2 vector. PCR products containing the FCR monomers were amplified using the pJET1.2 forward and reverse sequencing primers. 1µg of PCR product was mixed with 5µl of 25µM random hexamer primer and water was added up to 38µl. The PCR product was denatured at 95°C for 10 min and then snap cooled on ice. During denaturation 2.5µl of 1mM dA,C,G, 2.5µl of 1mM Alexa fluorophore conjugated dUTP, 5µl of 10X NEB Buffer 2, and 2µl of Klenow (exo-) polymerase were premixed. Both Alexa 488 and 568 dUTP conjugated fluorophores were used in these experiments. Nucleotide and polymerase mix was then mixed with denatured template and primers and incubated, protected from light, at 37°C overnight. The reaction was quenched with 2µl of 10mM EDTA and desalted with Microbio-spin 6 column (BioRad) to remove unincorporated nucleotides. Probes were then precipitated by adding 10µl of 10mg/ml salmon sperm DNA, 6µl of 3M Sodium Acetate, and 120µl 100% EtOH. Probes were vortexed and precipitated at -80°C for at least 30min. Samples were spun ∼16,000g at 4°C for 10min to pellet probes. Supernatant was removed and the pellet was resuspended in 1ml 70% EtOH, and spun again to wash. Supernatant was removed again and the pellet was allowed to air dry. Probes were then resuspended in 50µl hybridization buffer (65% Formamide, 5X SSC, 5X Denhardt’s Buffer (0.1% Ficoll-400, 0.1% Polyvinylpyrrolidone), with 150µg/ml yeast tRNA, and 0.5mg/ml salmon sperm DNA). Probes were incubated at 37°C for 15 min to allow complete solubilization and stored at -20°C for two to three months. When used for single-color FISH experiments 4µl of probe was mixed with 4µl of hybridization buffer, and for two-color FISH experiments 4µl of each probe were diluted together.

### Phylogram generation

All CENP-A associated sequences with at least 20 enriched k-mers were isolated. Enriched k-mers were defined as those with an centromere enrichment score above 25 median absolute deviations away from the mean. These sequences were then entered into sequential rounds of cluster generation based on sequence similarity using cd-hit-est, first clustering sequences together that were 98% identical, then 95%, and finally 90% identical by sequence (Fu et al. 2012). This generated a list of representative sequences. The top 50 most abundant sequences were then used to generate the phylogram using Genious Tree Builder with the following settings: Genetic Distance Model=Tamura-Nei, Tree building method=Neighbor-joining, Outgroup=No outgroup, Alignment Type=Global alignment, Cost Matrix=93% similarity. Colors were manually added to branches that contain FCR monomers used for validation by FISH. The originally identified Fcr1 monomer is labelled with an arrow. GC content of top 50 FCR monomers and input reads were plotted using FASTQC (Andrews and Others 2010).

### FISH data analysis Probe correlations and heatmap generation

Axial projections of images were analyzed as previously described (Moree 2011). CENP-C immunofluorescence signal was used as the centromere fiducial marker. Image specific background values were subtracted from FISH intensities at each centromere. For quantification of percentage of centromeres per nucleus positive for a FISH probe a cutoff background subtracted intensity of 200AU was used to call a centromere positive for negative. A custom bash script was used to quantify the percentage of centromeres per nucleus that were FISH positive. Quantifications for single color fish are from three or four experimental replicates with at 10-12 nuclei quantified per experiment. For two color FISH experiments background subtracted centromere intensities were plotted on a scatterplot and Pearson correlations were calculated in R. Two color FISH experiments were performed in duplicate with at least 200 centromeres quantified per slide per experiment. Results shown are from one representative experiment.

### Genome segment analysis

In order to identify regions on each chromosome that contain CENP-A enriched k-mers, an updated version of the *X. laevis* genome was first separated into 50kb segments using a custom python script. The bbduk program was then used to extract genome segments that had enriched k-mers. Enriched k-mers were defined by the ratio of normalized k-mer abundance in the CENP-A and INPUT libraries. Cutoffs were established by variable multiples of median absolute deviation from the median enrichment ratio. A stringent cutoff of 17 median absolute deviations from the median enrichment ration of all k-mers was chosen for downstream analysis. Genome segments with a minimal k-mer density were used to characterize centromeres on each chromosome by its k-mer content. 84 total 50kb genome segments were identified as containing a high abundance of CENP-A enriched k-mers. A binary matrix was generated indicating the presence or absence of each enriched k-mer on each genome segment. This matrix was then plotted as a heatmap clustering both axes (method= “jaccard”) to show both k-mers and genome segments that are found together or without clustering by genome segment to preserve the ordering of source chromosomes from each segment. Additionally chromosomes were clustered by the abundance of each CENP-A enriched k-mer (method= “euclidean”), after collapsing 50kb genome segments by chromosome. For genome visualization CENP-A enriched k-mers and genome segments with CENP-A enriched k-mers were aligned to the updated *X. laevis* genome using bowtie2 (Ben Langmead and Salzberg 2012) allowing for multiple alignment of k-mers. Genome alignments were then visualized using the pygenometracks python module (Ramírez et al. 2018). To estimate repeat array size, bedgraphs with 1bp resolution were created using deepTools2 (Ramírez et al. 2016). From this bedgraph the centromeric region on each chromosome was selected manually based on the 50kb genome segment analysis and then the centromere length was defined as the distance between the first and last base pair that had a CENP-A enriched k-mer aligned. We also report the total base pairs within the centromere on each chromosome that have CENP-A enriched k-mers aligned, and the fraction of the centromere with a CENP-A enriched kmer.

### Bowtie analysis

FCR monomers (150bp) were initially split into 25bp k-mers (126 total) using KMC (Kokot, Długosz, and Deorowicz 2017). K-mers from each monomer were then aligned to the *Xenopus laevis* genome 10.2 provided by Jessen Bredeson and Dan Rokhsar using bowtie1 (B. Langmead et al. 2009) allowing for no mismatches and for alignment as many times as possible. Alignment files were then used to count the number of times k-mers from each FCR monomer aligned to each chromosome. This produced a table of counts of the number of times k-mers from each FCR monomer aligned to each chromosome. This table was then plotted as a heatmap, normalizing the intensity to the highest count on each chromosome. This same analysis was also performed with k-mers that were specific to individual FCR monomers. Alignment counts were normalized by the number of k-mers that were specific to each FCR monomer to account for differences in the number of specific k-mers for each FCR monomer.

### Data Access

All raw and processed sequencing data generated in this study have been submitted to the NCBI Gene Expression Omnibus (GEO; https://www.ncbi.nlm.nih.gov/geo/) under accession number GSE153058. Our k-mer analysis pipeline is available at https://github.com/straightlab/xenla-cen-dna-paper.

## Acknowledgements

This work was supported by NIH R01 GM074728 to AFS and NIH R35 GM 118183 to RMH. OKS was supported by NIH T32 GM113854-02 and an NSF GRFP. We thank Magdalena Strzelecka and Andrew Grenfell for discussion. We thank members of the Straight laboratory for comments on the manuscript. We thank Jessen Bredeson and Daniel Rokhsar for providing early access to *X. laevis* genome release 10.2.

## References

Akiyoshi, Bungo, Krishna K. Sarangapani, Andrew F. Powers, Christian R. Nelson, Steve L. Reichow, Hugo Arellano-Santoyo, Tamir Gonen, Jeffrey A. Ranish, Charles L. Asbury, and Sue Biggins. 2010. “Tension Directly Stabilizes Reconstituted Kinetochore-Microtubule Attachments.” Nature. https://doi.org/10.1038/nature09594.

Andrews, Simon, and Others. 2010. “FastQC: A Quality Control Tool for High Throughput Sequence Data.” Babraham Bioinformatics, Babraham Institute, Cambridge, United Kingdom.

Desai, A., H. W. Deacon, C. E. Walczak, and T. J. Mitchison. 1997. “A Method That Allows the Assembly of Kinetochore Components onto Chromosomes Condensed in Clarified Xenopus Egg Extracts.” Proceedings of the National Academy of Sciences of the United States of America 94 (23): 12378–83.

Edwards, N. S., and A. W. Murray. 2005. “Identification of Xenopus CENP-A and an Associated Centromeric DNA Repeat.” Molecular Biology of the Cell 16 (4): 1800–1810.

Foley, E. A., and T. M. Kapoor. 2013. “Microtubule Attachment and Spindle Assembly Checkpoint Signalling at the Kinetochore.” Nature Reviews. Molecular Cell Biology 14 (1): 25–37.

Fu, L., B. Niu, Z. Zhu, S. Wu, and W. Li. 2012. “CD-HIT: Accelerated for Clustering the next-Generation Sequencing Data.” Bioinformatics 28 (23): 3150–52.

Guse, A., C. W. Carroll, B. Moree, C. J. Fuller, and A. F. Straight. 2011. “In Vitro Centromere and Kinetochore Assembly on Defined Chromatin Templates.” Nature 477 (7364): 354–58.

Harrington, J. J., G. Van Bokkelen, R. W. Mays, K. Gustashaw, and H. F. Willard. 1997. “Formation of de Novo Centromeres and Construction of First-Generation Human Artificial Microchromosomes.” Nature Genetics 15 (4): 345–55.

Hayden, K. E., E. D. Strome, S. L. Merrett, H. R. Lee, M. K. Rudd, and H. F. Willard. 2013. “Sequences Associated with Centromere Competency in the Human Genome.” Molecular and Cellular Biology 33 (4): 763–72.

Hayden, K. E., and H. F. Willard. 2012. “Composition and Organization of Active Centromere Sequences in Complex Genomes.” BMC Genomics 13. https://doi.org/ Artn 324 10.1186/1471-2164-13-324.

Hyman, A. A., K. Middleton, M. Centola, T. J. Mitchison, and J. Carbon. 1992. “Microtubule-Motor Activity of a Yeast Centromere-Binding Protein Complex.” Nature 359 (6395): 533–36.

Kasinathan, Sivakanthan, and Steven Henikoff. 2018. “Non-B-Form DNA Is Enriched at Centromeres.” Molecular Biology and Evolution 35 (4): 949–62.

Kokot, Marek, Maciej Długosz, and Sebastian Deorowicz. 2017. “KMC 3: Counting and Manipulating K-Mer Statistics.” Bioinformatics. https://doi.org/10.1093/bioinformatics/btx304.

Langmead, Ben, and Steven L. Salzberg. 2012. “Fast Gapped-Read Alignment with Bowtie 2.” Nature Methods 9 (4): 357–59.

Langmead, B., C. Trapnell, M. Pop, and S. L. Salzberg. 2009. “Ultrafast and Memory-Efficient Alignment of Short DNA Sequences to the Human Genome.” Genome Biology 10 (3): R25.

Logsdon, G. A., C. W. Gambogi, M. A. Liskovykh, E. J. Barrey, V. Larionov, K. H. Miga, P. Heun, and B. E. Black. 2019. “Human Artificial Chromosomes That Bypass Centromeric DNA.” Cell 178 (3): 624–39 e19.

Manuelidis, L. 1978. “Complex and Simple Sequences in Human Repeated DNAs.” Chromosoma 66 (1): 1–21.

McDermid, H. E., A. M. Duncan, M. J. Higgins, J. L. Hamerton, E. Rector, K. R. Brasch, and B. N. White. 1986. “Isolation and Characterization of an Alpha-Satellite Repeated Sequence from Human Chromosome 22.” Chromosoma 94 (3): 228–34.

McNulty, S. M., and B. A. Sullivan. 2018. “Alpha Satellite DNA Biology: Finding Function in the Recesses of the Genome.” Chromosome Research: An International Journal on the Molecular, Supramolecular and Evolutionary Aspects of Chromosome Biology 26 (3): 115–38.

Melters, D. P., K. R. Bradnam, H. A. Young, N. Telis, M. R. May, J. G. Ruby, R. Sebra, et al. 2013. “Comparative Analysis of Tandem Repeats from Hundreds of Species Reveals Unique Insights into Centromere Evolution.” Genome Biology 14 (1): R10.

Miga, K. H., Y. Newton, M. Jain, N. Altemose, H. F. Willard, and W. J. Kent. 2014. “Centromere Reference Models for Human Chromosomes X and Y Satellite Arrays.” Genome Research 24 (4): 697–707.

Moree, B., C. B. Meyer, C. J. Fuller, and A. F. Straight. 2011. “CENP-C Recruits M18BP1 to Centromeres to Promote CENP-A Chromatin Assembly.” The Journal of Cell Biology 194 (6): 855–71.

Musacchio, A., and A. Desai. 2017. “A Molecular View of Kinetochore Assembly and Function.” Biology 6 (1). https://doi.org/10.3390/biology6010005.

Ngan, V. K., and L. Clarke. 1997. “The Centromere Enhancer Mediates Centromere Activation in Schizosaccharomyces Pombe.” Molecular and Cellular Biology 17 (6): 3305–14.

Ng, R., and J. Carbon. 1987. “Mutational and in Vitro Protein-Binding Studies on Centromere DNA from Saccharomyces Cerevisiae.” Molecular and Cellular Biology 7 (12): 4522–34.

Ohzeki, J., V. Larionov, W. C. Earnshaw, and H. Masumoto. 2015. “Genetic and Epigenetic Regulation of Centromeres: A Look at HAC Formation.” Chromosome Research: An International Journal on the Molecular, Supramolecular and Evolutionary Aspects of Chromosome Biology 23 (1): 87–103.

Ohzeki, J., M. Nakano, T. Okada, and H. Masumoto. 2002. “CENP-B Box Is Required for de Novo Centromere Chromatin Assembly on Human Alphoid DNA.” The Journal of Cell Biology 159 (5): 765–75.

Peacock, W. J., D. Brutlag, E. Goldring, R. Appels, C. W. Hinton, and D. L. Lindsley. 1974. “The Organization of Highly Repeated DNA Sequences in Drosophila Melanogaster Chromosomes.” Cold Spring Harbor Symposia on Quantitative Biology 38: 405–16.

Ramírez, Fidel, Vivek Bhardwaj, Laura Arrigoni, Kin Chung Lam, Björn A. Grüning, José Villaveces, Bianca Habermann, Asifa Akhtar, and Thomas Manke. 2018. “High-Resolution TADs Reveal DNA Sequences Underlying Genome Organization in Flies.” Nature Communications. https://doi.org/10.1038/s41467-017-02525-w.

Ramírez, Fidel, Devon P. Ryan, Björn Grüning, Vivek Bhardwaj, Fabian Kilpert, Andreas S. Richter, Steffen Heyne, Friederike Dündar, and Thomas Manke. 2016. “deepTools2: A next Generation Web Server for Deep-Sequencing Data Analysis.” Nucleic Acids Research 44 (W1): W160–65.

Rudd, M. K., M. G. Schueler, and H. F. Willard. 2003. “Sequence Organization and Functional Annotation of Human Centromeres.” Cold Spring Harbor Symposia on Quantitative Biology 68: 141–49.

Session, A. M., Y. Uno, T. Kwon, J. A. Chapman, A. Toyoda, S. Takahashi, A. Fukui, et al. 2016. “Genome Evolution in the Allotetraploid Frog Xenopus Laevis.” Nature 538 (7625): 336–43.

Sorger, P. K., F. F. Severin, and A. A. Hyman. 1994. “Factors Required for the Binding of Reassembled Yeast Kinetochores to Microtubules in Vitro.” The Journal of Cell Biology. https://doi.org/10.1083/jcb.127.4.995.

Sullivan, Lori L., Kimberline Chew, and Beth A. Sullivan. 2017. “α Satellite DNA Variation and Function of the Human Centromere.” Nucleus 8 (4): 331–39.

Sun, Xiaoping, Hiep D. Le, Janice M. Wahlstrom, and Gary H. Karpen. 2003. “Sequence Analysis of a Functional Drosophila Centromere.” Genome Research 13 (2): 182–94.

Sun, X., J. Wahlstrom, and G. Karpen. 1997. “Molecular Structure of a Functional Drosophila Centromere.” Cell 91 (7): 1007–19.

Willard, H. F., and J. S. Waye. 1987. “Hierarchical Order in Chromosome-Specific Human Alpha-Satellite DNA.” Trends in Genetics: TIG 3 (7): 192–98.

Zasadzinska, Ewelina, and Daniel R. Foltz. 2017. “Orchestrating the Specific Assembly of Centromeric Nucleosomes.” Progress in Molecular and Subcellular Biology 56: 165–92.

